# Effects of various carbohydrates on the in vitro pollen germination of *Vinca rosea* and *Cucumis melo var. utilissimus*

**DOI:** 10.1101/2020.10.03.316216

**Authors:** S Sharma, A D Marni, V Sharma, P Manchanda

## Abstract

Pollen germination is crucial for the process of plant development which strongly depends on the presence of carbohydrates as a primary source of energy. In this study, we analyzed the differential effects of four sugars with varying concentrations on the pollen germination of *Vinca rosea Linn*. and *Cucumis melo var. utilissimus* (Duth. & Fuller) using Brewbaker and Kwack’s medium as germination medium and hanging drop method after an incubation period of 1 hour. Sucrose and glucose supported the pollen germination but galactose and fructose did not support and even considerably inhibited the pollen germination of *Vinca rosea*. However in pollen germination of *Cucumis melo var. utilissimus*, all the four sugars supported the pollen germination. The study suggests that 15% sucrose, for *Vinca rosea*, and 12% galactose, for *Cucumis melo var. utilissimus*, supports in achieving the highest pollen germination percentage when added to the pollen germination medium.

## Introduction

The development of pollen grains on stigma or in the in vitro conditions is a time-bound and high energy requiring process ^15^. The recognition of compatible pollen and the growth of pollen tube largely depends upon the receptivity of stigma and other recognition factors. The further growth of pollen tube, till it reaches embryo sac, is facilitated by the environment of stylar canal ^2^. Carbohydrates, act as an osmotic regulator and also as a substrate for primary energy source, supporting pollen germination and the pollen tube growth in stylar canal ^12^.

Most of the studies conducted on *Vinca rosea* are based on its therapeutic properties. In Ayurveda, extracts of both roots and shoots are used to treat several diseases. Many *vinca* alkaloids including vinblastine and vincristine are used for treating leukemia and Hodgkin’s lymphoma ^8^. *Cucumis melo var. utilissimus* is commonly called as American cucumber or “kakri” in India. The fruits are used for its juice or made into powder for cooking ^7^. Extract from the fruits is used to promote skin hydration, treat light burns and act as a stomach tonic. Seeds are used for suppressing cough, reducing fever and act as digestive aid ^4^. The present study investigates the effects of various sugars with varying concentrations on the in vitro pollen germination of *Vinca rosea* and *Cucumis melo var. utilissimus* by using a pollen germination medium and calculating pollen germination percentage. Different effects of sugars and their respective concentrations were observed on the pollen germination of Vinca rosea and Cucumis melo var. utilissimus. 15% sucrose and 12% galactose were found to be the best sugar concentrations along with the pollen germination medium to achieve the highest pollen germination percentage after an incubation period of 1 hour for *Vinca rosea* and *Cucumis melo var. utilissimus* respectively.

## Materials and Methods

### Pollen collection

Flowers of *Vinca rosea* and *cucumis melo var. utilissimus*, at anthesis, were collected from the botanical garden of Miranda House, University of Delhi during daytime. Pollen grains were carefully collected on separate clean petri dishes by tapping and brushing the anthers of each flower.

### Preparation of Pollen Germination Medium

Standard solution (100ml) of Brewbaker and Kwack’s medium ^7^ was prepared by using 0.01g of boric acid, 0.01g of potassium nitrate, 0.02g of magnesium sulphate heptahydrate and 0.03g of calcium nitrate. The medium was prepared by using four different sugars: sucrose, glucose, fructose and galactose. For each sugar, four different concentrations were made: 5%, 10%, 12% and 15%. The germination medium was autoclaved to maintain the sterility.

### Experimental Set-up

Hanging drop method ^16^ with slight modifications was used for performing the in vitro pollen germination. A small drop of pollen germination medium was taken on a glass slide and the pollen grains were transferred onto it with the aid of a nylon brush. The glass slide was inverted and placed on a glass cavity block. The edges were sealed with the help of petroleum jelly. The set-up was incubated in light with temperature set at 30°C.

### Observation of pollen germination

Observations for pollen germination was taken by microscopy four times, each with 15 mins interval. Three microscopic field views per replicate, that contained a minimum of 30 pollen grains, was observed. Pollen grains were considered germinated when the length of the pollen tube was greater than the diameter of the pollen grain. The total number of pollen grains that germinated was calculated and the percentage germination was calculated by dividing the number of pollen grains germinated by the total number of pollen grains.

### Statistical analysis

Three biological replicates for the experiment were performed using the above methodology. Descriptive statistics was used to determine mean and standard deviation for pollen germination percentage. Duncan’s multiple range test (DMRT) was used to determine the significant differences in pollen germination percentage among different sugar concentrations.

## Results

### Pollen germination in Vinca rosea

The pollen germination percentage for Vinca rosea under the effect of various sugars with varying concentrations was obtained as shown in Table 1. As presented, pollen grains germinated in every media except when fructose was used (Fig. 1a.-1d.). 5% sucrose concentration had an inhibitory effect on pollen germination. Using 15% sucrose in germination medium resulted in highest pollen germination percentage (67.99%) while the lowest pollen germination (3.06%) was obtained by using 10% galactose in germination medium after an incubation period of an hour. Lower concentrations of both glucose and galactose had an inhibitory effect on the pollen germination of *Vinca rosea*. Fructose, in any concentration, inhibited the pollen germination.

**Table 1.** Effect of various sugars with varying concentration on the pollen germination of *Vinca rosea*. Values (in %) are represented as Mean ± S.D. Within columns values of each parameter with the same superscript are not significantly different at (*P* = 0.05) Duncan’s multiple range test (DMRT).

| Time (min) | Sucrose Concentration |  |  |  |  |
| --- | --- | --- | --- | --- | --- |
|  | 5% | 10% | 12% | 15% | Control |
| 15 | 0 ± 0a | 0 ± 0a | 0 ± 0a | 25.7936 ± 0.8361a | 1.6269 ± 1.4102a |
| 30 | 0 ± 0a | 2.0634 ± 1.8028b | 3.4555 ± 1.2370b | 31.7272 ± 0.5662b | 1.7105 ± 1.4827a |
| 45 | 0 ± 0a | 9.1547 ± 0.7364c | 15.1532 ± 0.7780c | 42.1530 ± 0.6243c | 3.5776 ± 1.2426b |
| 60 | 0 ± 0a | 12.5216 ± 1.1040d | 23.2478 ± 0.4986d | 67.996 ± 1.1547d | 4.4014 ± 1.2714c |
| Time (min) | Glucose Concentration |  |  |  |  |
|  | 5% | 10% | 12% | 15% | Control |
| 15 | 0 ± 0a | 0 ± 0a | 0 ± 0a | 0 ± 0a | 1.6269 ± 1.4102a |
| 30 | 0 ± 0a | 0 ± 0a | 0 ± 0a | 7.4853 ± 0.7090b | 1.7105 ± 1.4827a |
| 45 | 0 ± 0a | 0 ± 0a | 0 ± 0a | 10.4323 ± 0.9278c | 3.5776 ± 1.2426b |
| 60 | 0 ± 0a | 0 ± 0a | 0 ± 0a | 23.4508 ± 0.8336d | 4.4014 ± 1.2714c |
| Time (min) | Galactose Concentration |  |  |  |  |
|  | 5% | 10% | 12% | 15% | Control |
| 15 | 0 ± 0a | 0 ± 0a | 0 ± 0a | 0 ± 0a | 1.6269 ± 1.4102a |
| 30 | 0 ± 0a | 0 ± 0a | 3.0786 ± 0.2806b | 0 ± 0a | 1.7105 ± 1.4827a |
| 45 | 0 ± 0a | 0 ± 0a | 5.6903 ± 0.4949c | 0 ± 0a | 3.5776 ± 1.2426b |
| 60 | 0 ± 0a | 3.0693 ± 0.1905b | 12.8835 ± 1.2903d | 0 ± 0a | 4.4014 ± 1.2714b |
| Time (min) | Fructose Concentration |  |  |  |  |
|  | 5% | 10% | 12% | 15% | Control |
| 15 | 0 ± 0a | 0 ± 0a | 0 ± 0a | 0 ± 0a | 1.6269 ± 1.4102a |
| 30 | 0 ± 0a | 0 ± 0a | 0 ± 0a | 0 ± 0a | 1.7105 ± 1.4827a |
| 45 | 0 ± 0a | 0 ± 0a | 0 ± 0a | 0 ± 0a | 3.5776 ± 1.2426b |
| 60 | 0 ± 0a | 0 ± 0a | 0 ± 0a | 0 ± 0a | 4.4014 ± 1.2714c |

**Fig. 1.**
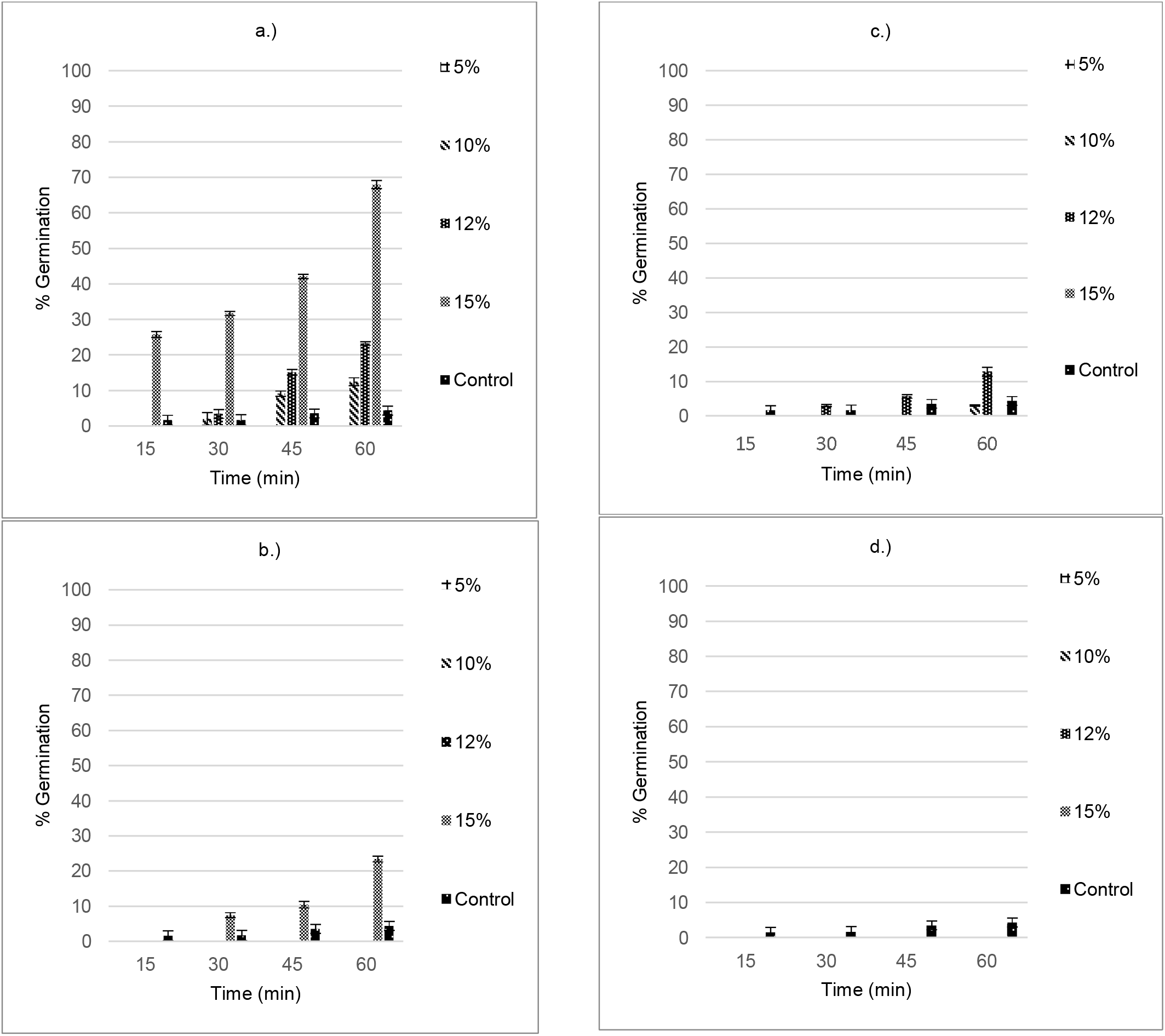
Effect of varying concentrations of a.) Sucrose, b.) Glucose, c.) Galactose and d.) Fructose on the in vitro pollen germination of Vinca rosea. Data are shown by Mean ± S.D.

### Pollen germination in *Cucumis melo* var. *utilissimus*

The pollen germination percentage for *Cucumis melo* var. *utilissimus* under the effect of various sugars with varying concentrations was obtained as shown in Table 2.

**Table 2.** Effect of various sugars with varying concentration on the pollen germination of *Cucumis melo*. Values (in %) are represented as Mean ± S.D. Within columns values of each parameter with the same superscript are not significantly different at (*P* = 0.05) Duncan’s multiple range test (DMRT).

| Time (min) | Sucrose Concentration |  |  |  |  |
| --- | --- | --- | --- | --- | --- |
|  | 5% | 10% | 12% | 15% | Control |
| 15 | 10.932 ± 0.6062a | 7.9194 ± 0.5337a | 16.4781 ± 0.7762a | 12.6515 ± 0.6200a | 2.7329 ± 0.1515a |
| 30 | 25.0221 ± 1.1519b | 12.4689 ± 0.9166b | 25.4757 ± 1.1330b | 20.3921 ± 0.3396b | 5.6198 ± 0.3202b |
| 45 | 33.9432 ± 0.5295c | 23.6438 ± 0.7673c | 36.8434 ± 0.5884c | 29.9636 ± 0.8164c | 10.0637 ± 1.1559c |
| 60 | 36.6987 ± 0.5306c | 28.0373 ± 0.8773d | 53.4214 ± 0.9768d | 34.2811 ± 0.9045d | 12.9308 ± 0.7233d |
| Time (min) | Glucose Concentration |  |  |  |  |
|  | 5% | 10% | 12% | 15% | Control |
| 15 | 15.6545 ± 0.2337a | 18.9507 ± 0.5462a | 9.0535 ± 0.8325a | 2.7834 ± 0.1548a | 2.7329 ± 0.1515a |
| 30 | 26.6025 ± 1.0612b | 30.2455 ± 0.7299b | 14.2483 ± 0.7924b | 9.5750 ± 1.0785b | 5.6198 ± 0.3202b |
| 45 | 32.4059 ± 0.0459c | 46.1841 ± 1.0849c | 21.1655 ± 1.0173c | 18.6612 ± 0.695c | 10.0637 ± 1.1559c |
| 60 | 41.7055 ± 0.7001d | 56.3911 ± 1.1185d | 31.9536 ± 0.6112d | 25.7282 ± 0.7353d | 12.9308 ± 0.7233d |
| Time (min) | Galactose Concentration |  |  |  |  |
|  | 5% | 10% | 12% | 15% | Control |
| 15 | 5.5814 ± 0.4667a | 13.7774 ± 0.5721a | 28.828 ± 0.6515a | 18.4244 ± 0.9229a | 2.7329 ± 0.1515a |
| 30 | 8.1438 ± 0.6624b | 21.5488 ± 0.5831b | 41.6864 ± 1.0166b | 24.5261 ± 1.1647b | 5.6198 ± 0.3202b |
| 45 | 12.6119 ± 0.4348c | 40.2146 ± 0.7107c | 53.5372 ± 0.8779c | 37.7841 ± 0.8211c | 10.0637 ± 1.1559c |
| 60 | 17.7137 ± 0.6764d | 54.8933 ± 0.6366d | 65.3399 ± 0.7144d | 48.9583 ± 1.8042d | 12.9308 ± 0.7233d |
| Time (min) | Fructose Concentration |  |  |  |  |
|  | 5% | 10% | 12% | 15% | Control |
| 15 | 11.6799 ± 0.7277a | 19.2885 ± 0.8009a | 5.8207 ± 0.6638a | 9.6757 ± 0.8514a | 2.7329 ± 0.1515a |
| 30 | 18.0022 ± 1.1831b | 33.3505 ± 0.9267b | 10.6516 ± 0.7224b | 12.7688 ± 0.2327b | 5.6198 ± 0.3202b |
| 45 | 22.87 ± 0.5724c | 46.9229 ± 0.5268c | 16.0279 ± 1.0168c | 18.1152 ± 0.5700c | 10.0637 ± 1.1559c |
| 60 | 35.7376 ± 0.6145d | 52.2535 ± 0.7847d | 25.9454 ± 0.3239d | 25.9726 ± 0.4390d | 12.9308 ± 0.7233d |

As depicted, every germination medium irrespective of the sugar and its concentration used, was successful in pollen germination (Fig. 2a.-2d.). Pollen grains showed the highest germination percentage (65.33%) when 12% galactose was used along with the germination medium and the lowest germination percentage (17.71%) was obtained when 5% galactose was used along with the germination medium after an incubation period of an hour.

**Fig. 2.**
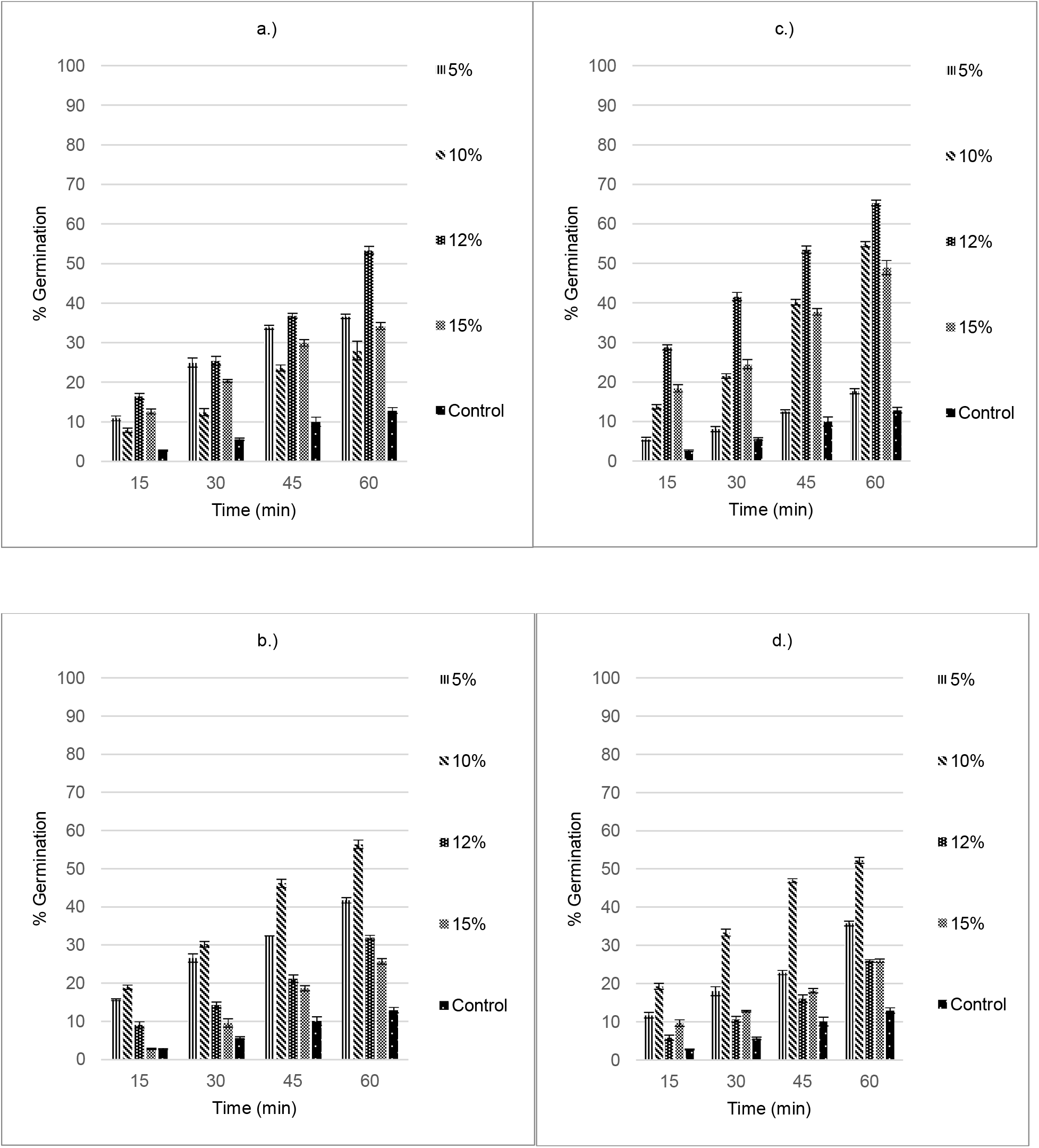
Effect of varying concentrations of a.) Sucrose, b.) Glucose, c.) Galactose and d.) Fructose on the in vitro pollen germination of Cucumis melo var. utilissimus. Data are shown by Mean ± S.D.

## Discussion

The present study demonstrates that the germination of *Vinca rosea* and *Cucumis melo* var. *utilissimus* pollen grains is largely influenced by the exogenous supply of various sugars with varying concentrations.

Numerous studies have shown that sucrose exhibits a strong stimulatory action on the growth of pollen grains ^10^. Both glucose and fructose have reported to show varied effects on pollen germination of various plant species ^9^. A higher concentration of sucrose as well as glucose promotes the in vitro pollen germination of *Vinca rosea* ^1^. Although, in case of *Vinca rosea*, the lower concentrations of glucose inhibit the pollen germination which supports the involvement of hexokinase mediated pathway of inhibiting sucrose mediated pollen germination ^5^. Sucrose promotes the pollen germination both in case of *Vinca rosea* and *Cucumis melo* var. *utilissimus* and this can be supported by the fact that sucrose is the most common form of sugar that is translocated to non-photosynthetic tissue, such as flower, and thus is also excreted by the stigma as well as the stylar canal for hydrating the pollen and supporting pollen tube growth ^6^. Glucose supports the pollen germination minimally in *Vinca rosea* as well as in *Cucumis melo* var. *utilissimus*. This could be due to the weak expression of AtSTP9, a glucose-specific monosaccharide transporter, in the mature pollen grain ^14^.

Fructose inhibits the pollen germination of *Vinca rosea* and not *Cucumis melo* var. *utilissimus*. Presence of fructose causes the pollen grains to completely utilize their stored sugars as respiration substrates and thus fail to develop ^9^. On the other hand, in case of *Cucumis melo* var. *utilissimus*, fructose promotes the pollen germination which can be supported by the studies which states the presence of fructose in the stigmatic fluid of plants promote pollen germination ^11^. Like that of fructose, the same effect can be clearly Seen with galactose which inhibits the pollen germination in Vinca rosea but largely promotes the pollen germination in Cucumis melo var. utilissimus. The inhibitory effect of galactose can be explained by the hexokinase mediated pathway of sucrose mediated pollen germination inhibition ^5^.

This study establishes the fact that the influence of various sugars on the germination of pollen grains also depends on the physiology of the pollen grains. This study adds to the information of developmental biology of pollen grains of *Vinca rosea*; which is a known ornamental and medicinal plant, and *Cucumis melo* var. *utilissimus*, which is a popular vegetable crop and medicinal plant of India. This study can further be used for extensive cultivation of these plants around the world.

## Acknowledgement

We thank the Department of Botany, Hindu College, University of Delhi for the lab materials and the equipment used in the study.

